## Supplementary File 1 for "Efflux pump gene amplifications bypass necessity of multiple target mutations for resistance against dual-targeting antibiotic"

**SUPPLEMENTAL FILE 1**

### Efflux pump gene amplifications bypass necessity of multiple target mutations for resistance against dual-targeting antibiotic

Kalinga Pavan T. Silva, Ganesh Sundar, Anupama Khare*

Laboratory of Molecular Biology, National Cancer Institute, National Institutes of Health, Bethesda, MD 20892, USA

#### SUPPLEMENTARY FIGURES AND FIGURE LEGENDS

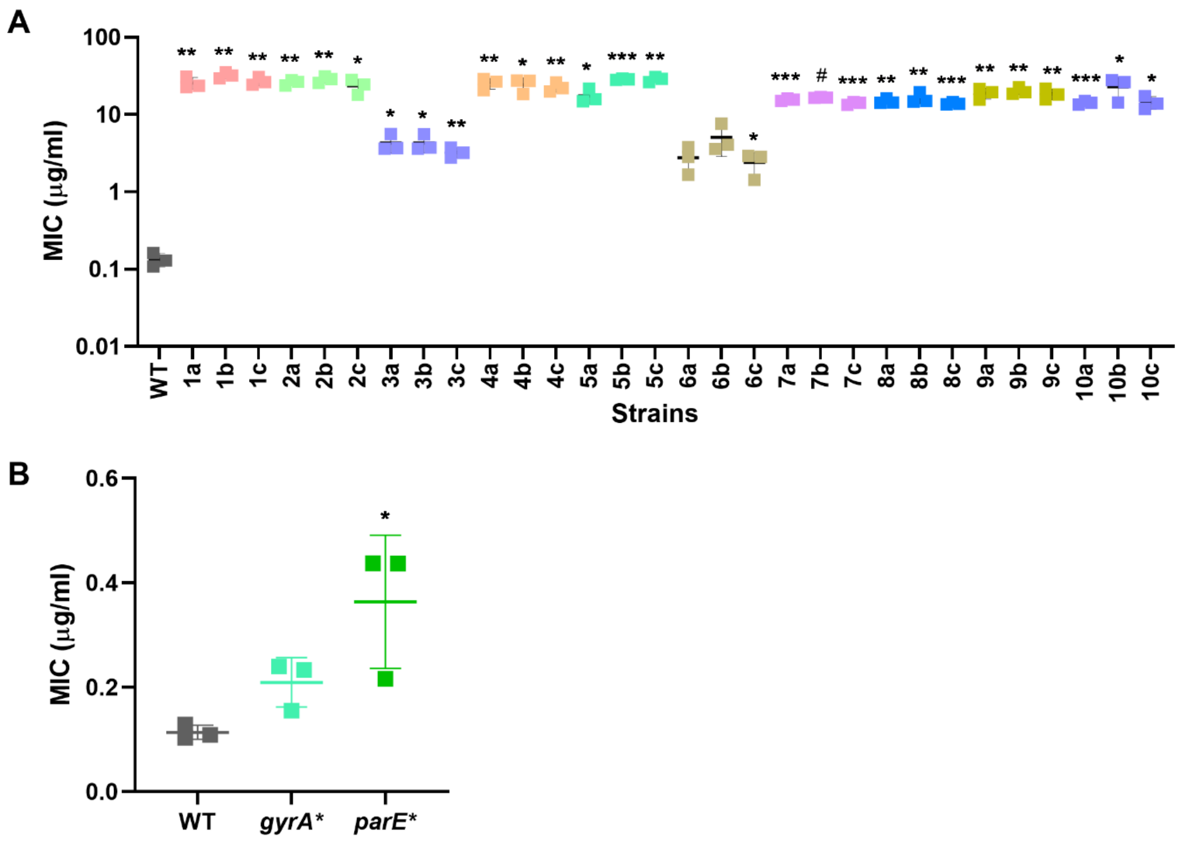

**Supplementary Figure 1. Evolved isolates are resistant to delafloxacin (DLX). (A)** DLX MICs of WT, and 3 isolates each from the final passage of all 10 independent populations, tested in MH2. **(B)** DLX MICs of WT, and allele-replacement strains carrying evolved alleles of *gyrA* (*gyrA*$^{E88K}$) and *parE* (*parE*$^{D432G}$) in MH2. Error bars show the standard deviation of three biological replicates. Significance for all is shown for comparison to the WT, as tested by **(A)** Brown-Forsythe and Welch ANOVA tests, followed by an unpaired t-test with Welch's correction for each comparison and **(B)** a one-way ANOVA with Holm-Sidak's test for multiple comparisons. (* $p < 0.05$, ** $p < 0.01$, *** $p < 0.001$, # $p < 0.0001$).

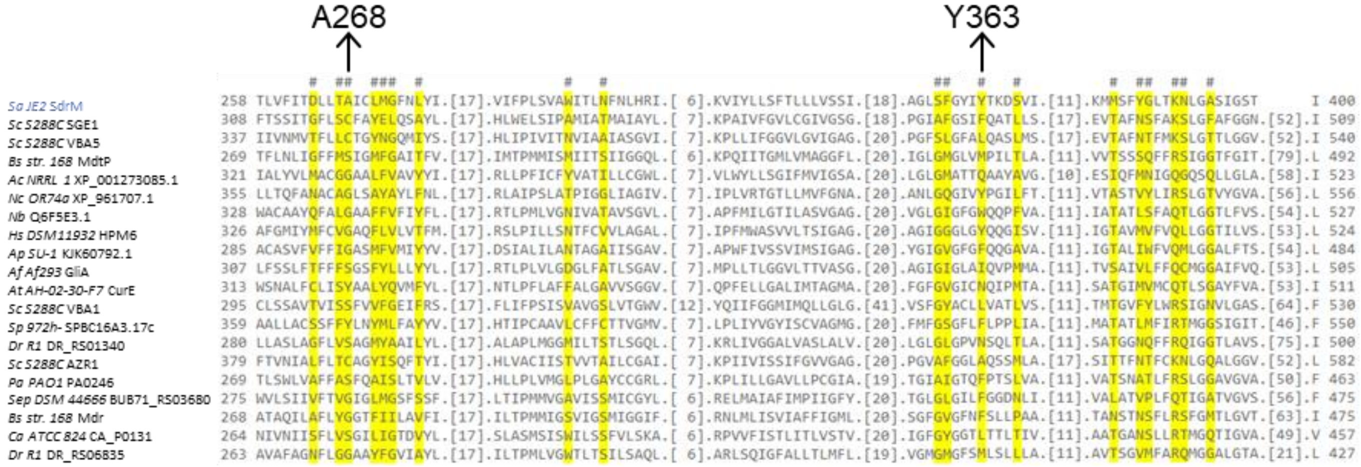

**Supplementary Figure 2. A268 and Y363 amino acid residues are predicted to be in the binding pocket of SdrM.** The SdrM amino acid sequence was aligned to similar MFS efflux pumps using the Conserved Domain Database (1) (https://www.ncbi.nlm.nih.gov/Structure/cdd/wrpsb.cgi?INPUT_TYPE=precalc&SEQUENCE=447160250). The amino acids predicted to be in the binding pocket are highlighted in yellow, and the SdrM A268 and Y363 residues are indicated. The other protein sequences are from the following species: *Sc*: *Saccharomyces cerevisiae*, *Bs*: *Bacillus subtilis*, *Ac*: *Aspergillus clavatus*, *Nc*: *Neurospora crassa*, *Nb*: *Neocamarosporium betae*, *Hs*: *Hypomyces subiculosus*, *Ap*: *Aspergillus parasiticus*, *Af*: *Aspergillus fumigatus*, *At*: *Aspergillus terreus*, *Sp*: *Saccharomyces pombe*, *Dr*: *Deinococcus radiodurans*, *Pa*: *Pseudomonas aeruginosa*, *Sep*: *Seinonella peptonophila*, *Ca*: *Clostridium acetobutylicum*.

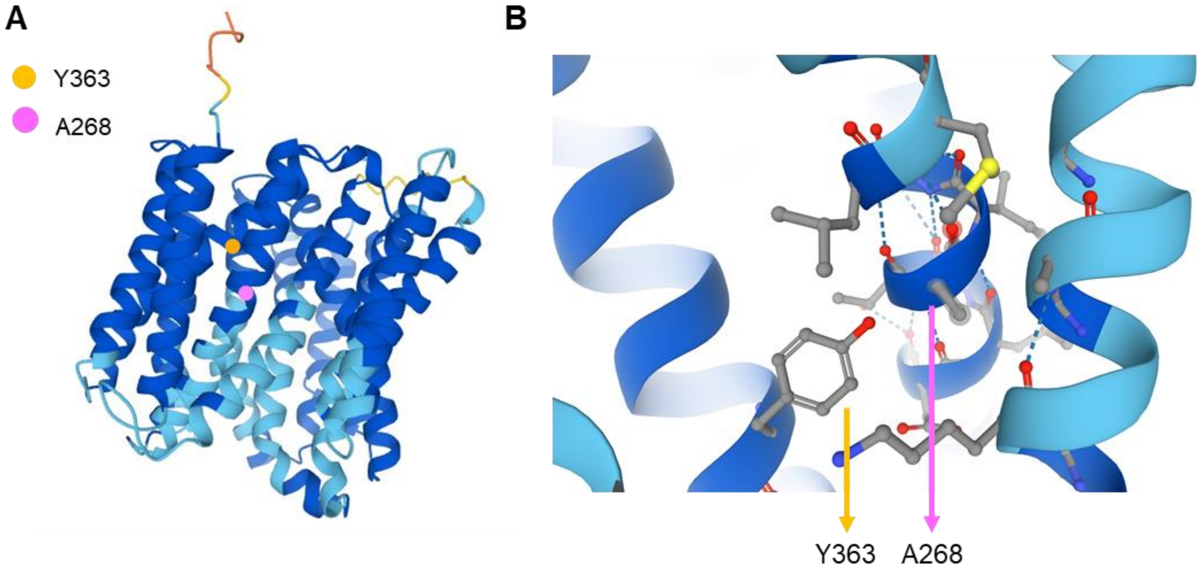

**Supplementary Figure 3. The A268 and Y363 residues are predicted to be in close proximity to each other.** AlphaFold (2, 3) prediction of **(A)** the entire SdrM structure, as well as **(B)** a zoomed-in region containing A268 and Y363.

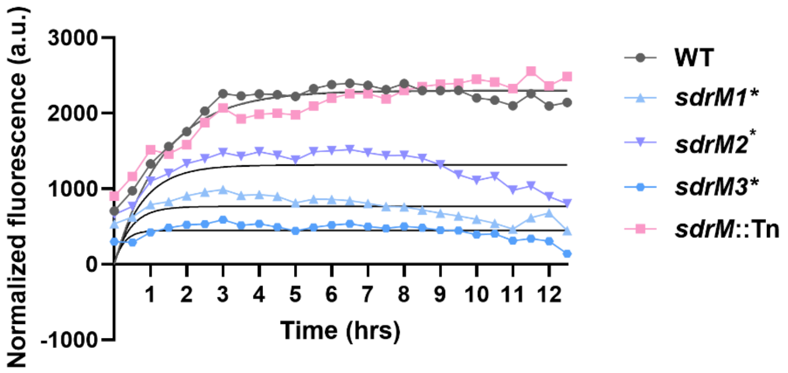

| | WT | *sdrM1** | *sdrM2** | *sdrM3** | *sdrM*::Tn |
|---|---|---|---|---|---|
| **Rate of efflux ($\rho_{out}$)** | 0.3647 | 1.833 | 0.9136 | 3.383 | 0.3666 |
| **95% confidence interval** | 0.3374 to 0.3941 | 1.628 to 2.085 | 0.8149 to 1.029 | 3.023 to 3.828 | 0.3294 to 0.4075 |

**Supplementary Figure 4. Modeled efflux rates of the allele-replacement strains are higher than the WT.** Normalized fluorescence (intrinsic DLX fluorescence/$OD_{600}$) was measured for the indicated strains (shown in **Figure 2B**). Least squares fit for the normalized fluorescence with rate of DLX efflux was determined. Shown are the mean values from three biological replicates with the lines representing the best fit curves. The best fit values for rate of efflux are indicated in the table, with the 95% confidence intervals.

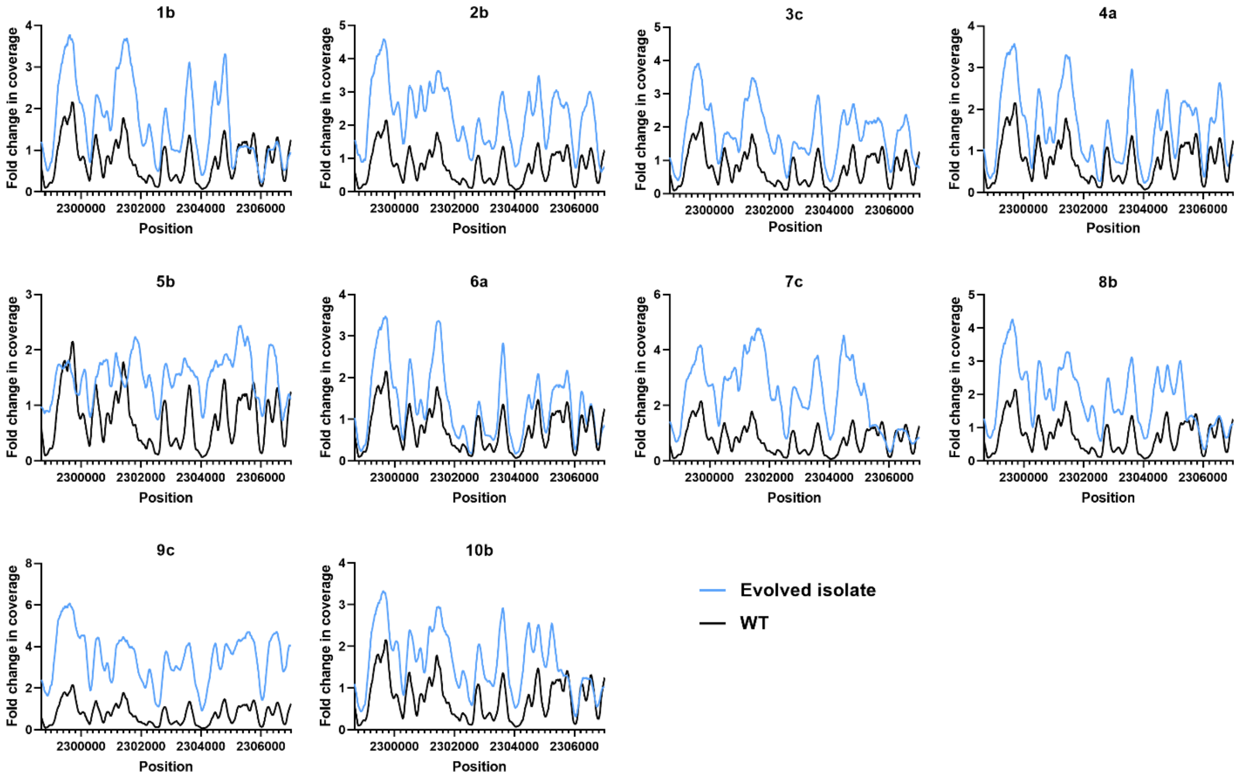

**Supplementary Figure 5. Efflux pump gene amplifications were seen in isolates from all evolved populations.** Relative read coverage of the amplified region compared to the entire genome shown for the WT, and one isolate from each evolved population. The lines represent a smoothed fit using a generalized additive model considering the nearest 100 neighboring nucleotides.

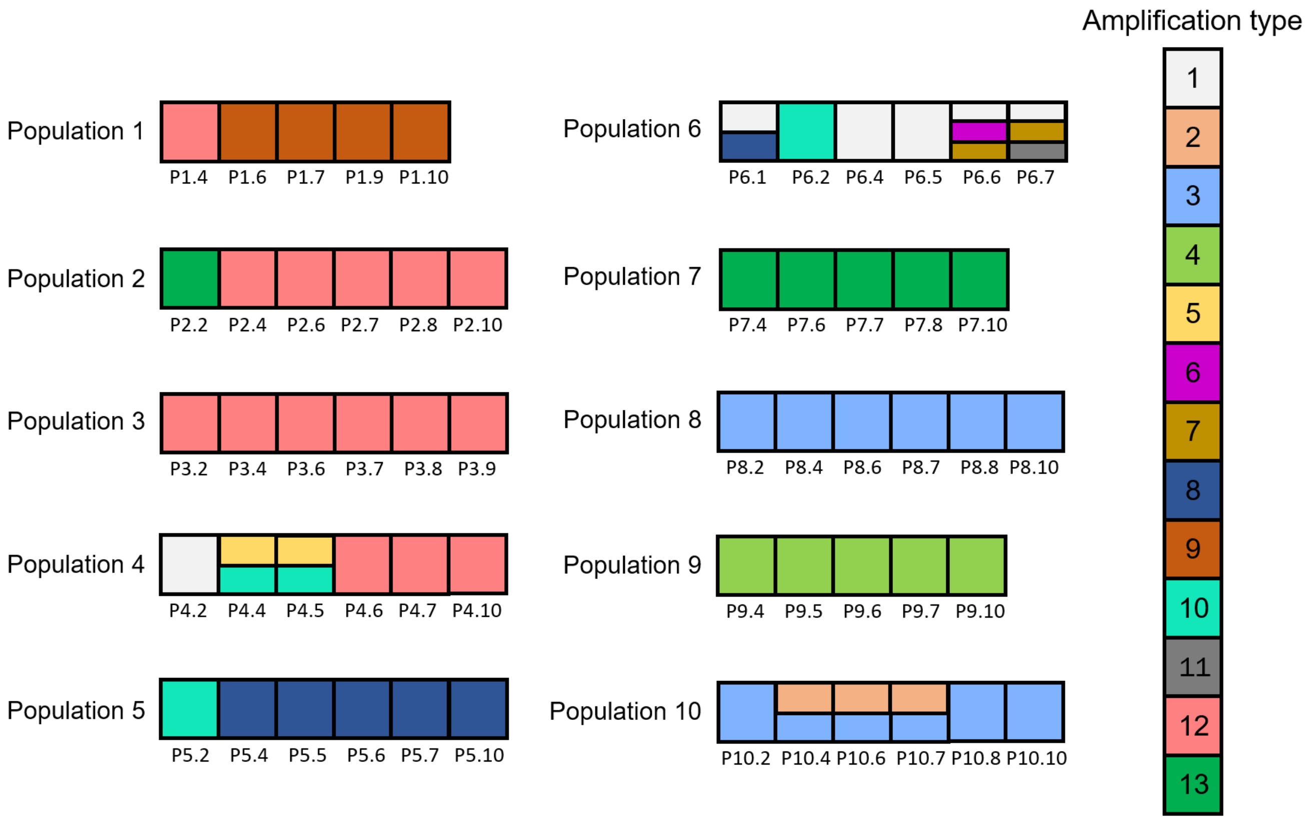

**Supplementary Figure 6. The amplifications of the *sdrM* genomic locus are dynamic within the evolving populations.** The amplification type(s) present in each sequenced passage from the ten independent populations are shown.

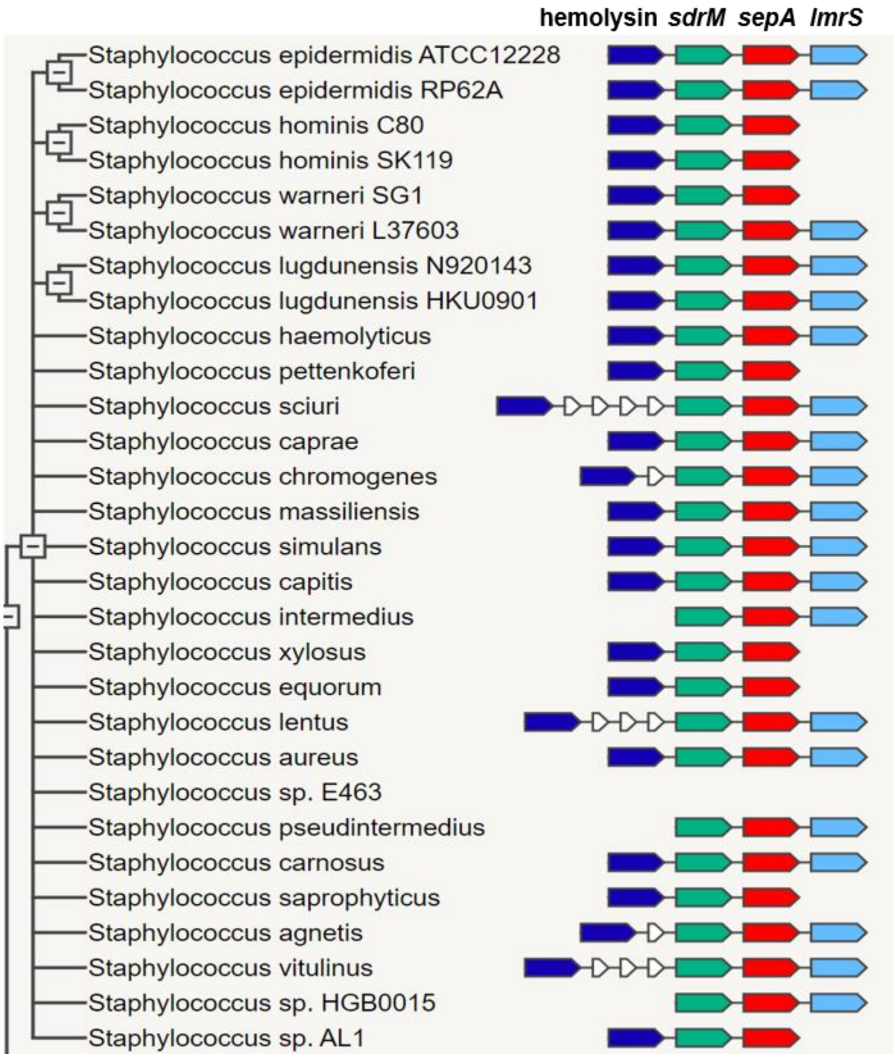

**Supplementary Figure 7. The *sdrM*, *sepA*, and *lmrS* genes are located adjacent to each other in most *Staphylococcus* species.** Gene neighborhood analysis using the STRING database (4) for genes located near *sdrM* is shown.

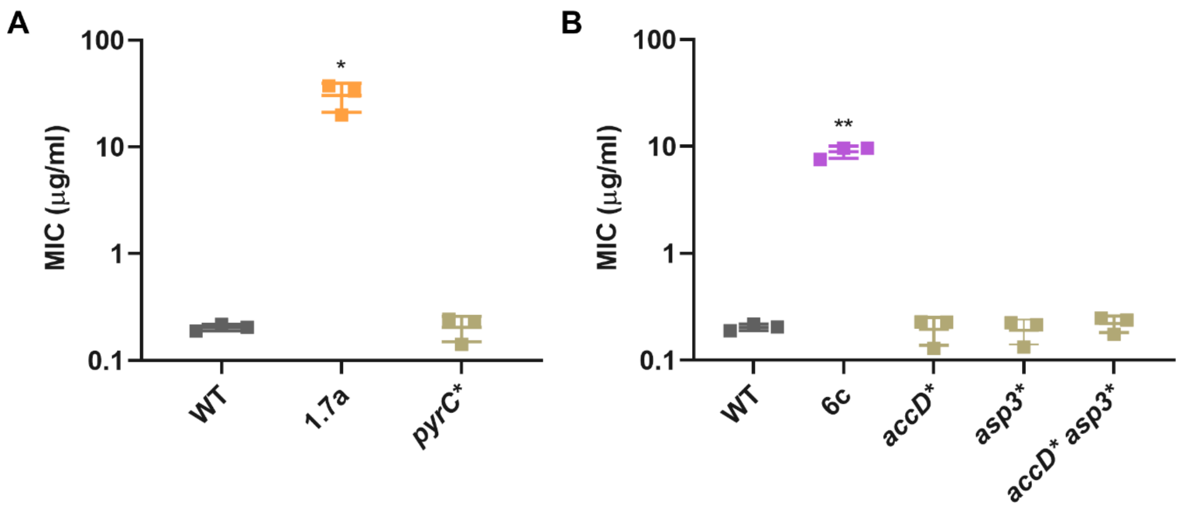

**Supplementary Figure 8. Additional mutations in evolved strains do not increase DLX resistance. (A, B).** The DLX MICs of WT, the evolved isolates 1.7a and 6c, and the indicated allelic replacement mutants were measured in M63. Error bars show the standard deviation of three biological replicates. Significance is indicated for comparison to the WT as tested by Brown-Forsythe and Welch ANOVA tests, followed by an unpaired t-test with Welch's correction for each comparison (* $p < 0.05$, ** $p < 0.01$).

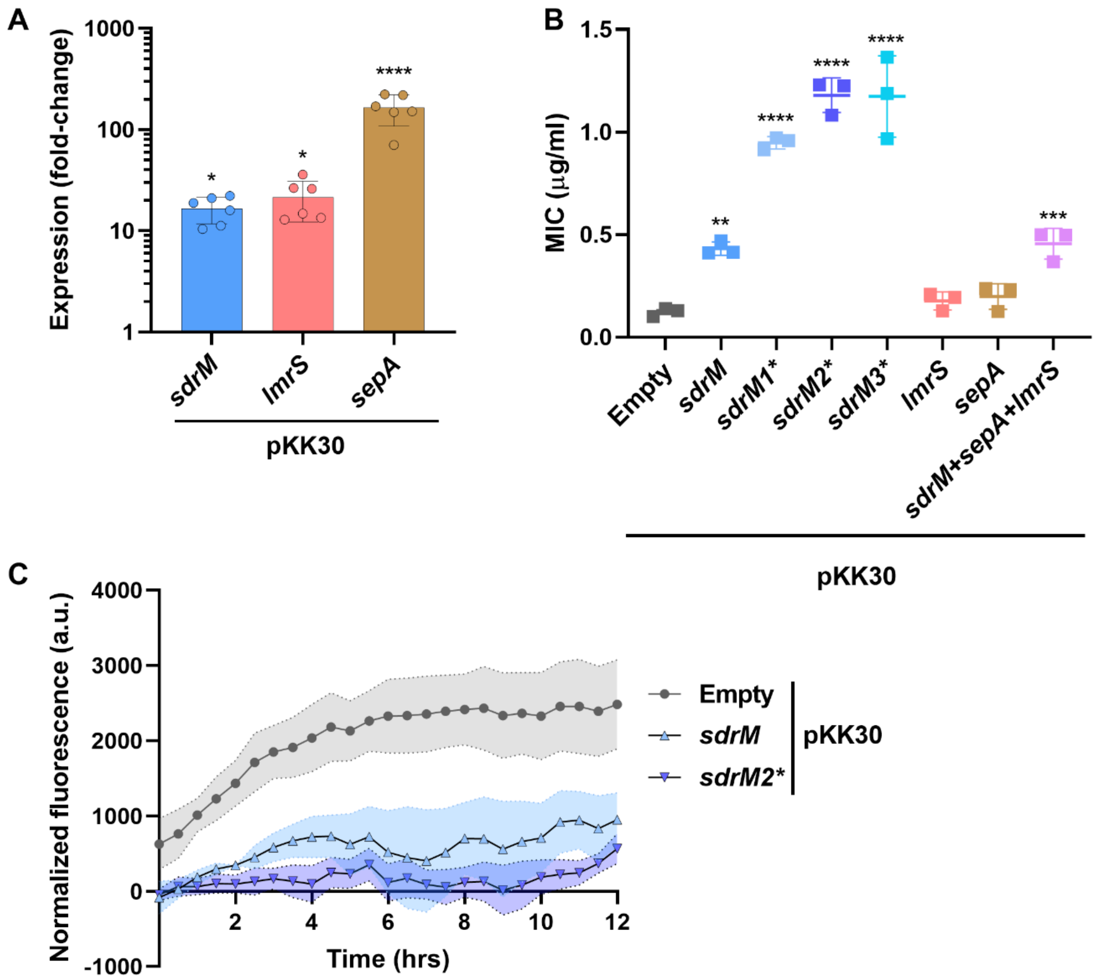

**Supplementary Figure 9. Overexpression of *sdrM* increases DLX resistance and efflux.** WT strains with either WT or mutant alleles of *sdrM*, *lmrS*, *sepA*, or all three WT efflux pumps expressed from pKK30 under their native promoters, were tested for expression, DLX resistance, and efflux. **(A)** Expression of the indicated pumps in the respective over-expression strains was measured by RT-qPCR. Data is shown as fold-change in expression compared to a WT strain carrying the pKK30 empty plasmid. Error bars show the standard deviation of three technical replicates each from two biological replicates. **(B)** DLX MICs were measured in M63. Error bars show the standard deviation of three biological replicates. **(C)** Normalized fluorescence (intrinsic fluorescence of DLX / $OD_{600}$) was measured for the indicated strains. Shaded areas represent the standard error of three biological replicates. Significance is shown for comparison to **(A)** a strain carrying the empty vector, set to 1, as tested by a Kruskal-Wallis test followed by an uncorrected Dunn's test for each comparison and **(B)** the strain with the empty pKK30 plasmid, as tested by a one-way ANOVA with Holm-Sidak's test for multiple comparisons (* $p < 0.05$, ** $p < 0.01$, *** $p < 0.001$, **** $p < 0.0001$).

#### SUPPLEMENTARY TABLES

**Supplementary Table 1. Evolution of DLX resistant populations and isolates.** (Excel)

**Supplementary Table 2. Canonical target and efflux pump mutations and amplifications present in the evolved populations and isolates.** (Excel)

**Supplementary Table 3. Sixteen different amplification types of *sdrM* seen in the evolved populations and isolates.** (Excel)

**Supplementary Table 4. Strains and plasmids used in this study.**

| Name | Description | Source |
|---|---|---|
| **Strains** | | |
| *S. aureus* | | |
| JE2 | *Staphylococcus aureus* subsp. *aureus* USA300_FPR3757 (CA-MRSA)-JE2 | (5) |
| RN4220 | Restriction modification deficient *S. aureus*; used to shuttle pKK30 derivatives into JE2 | (6-8) |
| SB229 | JE2 *sdrM1** | This study |
| SB225 | JE2 *sdrM2** | This study |
| SB262 | JE2 *sdrM3** | This study |
| SB267 | JE2 *gyrA** | This study |
| SB268 | JE2 *parE** | This study |
| SB236 | JE2 *pyrC** | This study |
| SB224 | JE2 *accD** | This study |
| SB234 | JE2 *asp3** | This study |
| SB235 | JE2 *accD* asp3** | This study |
| NE531 | JE2 *sdrM::*Tn | (9) |
| *E. coli* | | |
| DH5α-λpir | DH5α lysogenized with λpir; host for pKK30 | (10) |
| IM08B | For plasmid transfer into *S. aureus* | (11) |
| **Plasmids** | | |
| pIMAY* | *S. aureus* allelic exchange plasmid; $Cm^R$ | (12) |
| pKK30 | Expression vector for *S. aureus*; $Tmp^R$ | (13) |
| pSB271 | pKK30-*sdrM* | This study |
| pSB273 | pKK30-*sdrM1** | This study |
| pSB272 | pKK30-*sdrM2** | This study |
| pSB324 | pKK30-*sdrM3** | This study |
| pSB274 | pKK30-*lmrS* | This study |
| pSB275 | pKK30-*sepA* | This study |
| pSB414 | pKK30-*sdrM-sepA-lmrS* | This study |

$Cm^R$: chloramphenicol resistant; $Tmp^R$: trimethoprim resistant

**Supplementary Table 5. Primers used in this study.**

| Primer Description | Primer Sequence* |
|---|---|
| Amplify *sdrM* for pIMAY* F | GGTATCGATAAGCTTGATATCGAATTCGAAGGAGGTATTTCATGCGA |
| Amplify *sdrM* for pIMAY* R | ATTGGAGCTCCACCGCGGTGGCGGCCGCCGCACCAGAAAGTACAAAAA |
| Check pIMAY*-*sdrM* integration F | AATGAAAAGTCGCGCCTCTA |
| Check pIMAY*-*sdrM* integration R | AAAAGCTGGGGGAAACTCCG |
| Sanger sequencing for *sdrM1** and *sdrM2** | ATAAGCCCAATGCTAATTAC |
| Amplify *accD** for pIMAY* F | ATCGATAAGCTTGATATCGTGCGCTGTCACAAGATATG |
| Amplify *accD** for pIMAY* R | CTCCACCGCGGTGGCTCCACATCATTTTTATCTTGAGATTC |
| Check pIMAY*-*accD* integration F | TCTAGGGTCGTAGGTCTTTC |
| Check pIMAY*-*accD* integration R | GCAATTTTTGAAGGACGTTCATA |
| Sanger sequencing for *accD* | CAGCAACGCCAAATTTCATA |
| Amplify *asp3** for pIMAY* F | TGGAGCTCCACCGCGGTGGCCATTTCAAGAGCTATTACCTG |
| Amplify *asp3** for pIMAY* R | TATCGATAAGCTTGATATCGTTTAACTTCATCGCTCCATG |
| Check pIMAY*-*asp3* integration F | AGAAGAGTTCGGTACAGCATTAG |
| Check pIMAY*-*asp3* integration R | CTCGCTTCGCTAAATAATCATTC |
| Sanger sequencing for *asp3** | CGAGCCAAACATGACCAAAC |
| Amplify *pyrC** for pIMAY* F | TGGAGCTCCACCGCGGTGGCCGTAGTAATTACCATAGTTTAAAAGC |
| Amplify *pyrC** for pIMAY* R | TATCGATAAGCTTGATATCGCCTAAACGGTAGCCTTCG |
| Check pIMAY*-*pyrC* integration F | CCAATTGCGAATGCTGGTG |
| Check pIMAY*-*pyrC* integration R | GATCTGACCTGTATATGATGGATC |
| Sanger sequencing for *pyrC** | CCAATTGCGAATGCTGGTG |
| Amplify *sdrM3** for pIMAY* F | TGGAGCTCCACCGCGGTGGCTGATAGGCGGTGACTATG |
| Amplify *sdrM3** for pIMAY* R | TATCGATAAGCTTGATATCGCTAGTATCTAGGAAATTTATTATTTC |
| Check pIMAY*-*sdrM3* integration F and Sanger sequencing for *sdrM3** | TGTCACGTTACAAGTTCCTAGAG |
| Check pIMAY*-*sdrM3* integration R | CCATTAAATGCTCGACTGCAAATA |
| Amplify *gyrA* for pIMAY* F | TATCGATAAGCTTGATATCGCACGATACAAAGGTCTTG |
| Amplify *gyrA* for pIMAY* R | TGGAGCTCCACCGCGGTGGCTAGCATAAAAATAAGACTCCC |
| Check pIMAY*-*gyrA* integration F | TCTGAATTGAATCCAACACCAA |
| Check pIMAY*-*gyrA* integration R | GTGCCACCATCAAGACTTATCA |
| Sanger sequencing for *gyrA** | GAACTGAACTTTTGAAGGAG |
| Amplify *parE** for pIMAY* F | TATCGATAAGCTTGATATCGAGAATAACTATTGTATAGTTTTAAAAACG |
| Amplify *parE** for pIMAY* R | TGGAGCTCCACCGCGGTGGCATCACCTAAAACATCTTCAAG |
| Check pIMAY*-*parE* integration F | GGGAAAGCGCCGATAAGATA |
| Check pIMAY*-*parE* integration R | AGTCTCCATGTGGATGATATTG |

| | |
|---|---|
| Sanger sequencing for *parE** | TGAAGCTAGAAGTGCTGTTGAT |
| Gibson assembly for pKK30 F | GCGGCCGCTAGCCTAGGAGC |
| Gibson assembly for pKK30 R | ATCGCCTGTCACTTTGCTTGATATATGA |
| Amplify *sdrM* for pKK30 F | ATCAAGCAAAGTGACAGGCGATACAGTATTTATTTTATTATGGGGAAC |
| Amplify *sdrM* for pKK30 R | GAGCTCCTAGGCTAGCGGCCGCCTATTCTTTTGATTGAGATGAC |
| Amplify *sepA* for pKK30 F | ATCAAGCAAAGTGACAGGCGATAATGATGTCATTTTATGGATTAAC |
| Amplify *sepA* for pKK30 R | GAGCTCCTAGGCTAGCGGCCGCCTATTTTCTATTATTTAAATTTTTAC |
| Amplify *lmrS* for pKK30 F | ATCAAGCAAAGTGACAGGCGATTCAATATCGATTTTTGGGTC |
| Amplify *lmrS* for pKK30 R | GAGCTCCTAGGCTAGCGGCCGCTTAAAATTTCCTTCTATTACTTTC |
| Sanger sequencing for pKK30 F | CTGGGAAGTCGAATCTTCAGTAG |
| Sanger sequencing for pKK30 R | ATGATAGGTCTGGCAAAGCC |
| qPCR for *rpoC* F | CTGTGAAAGAATTTTCGGAC |
| qPCR for *rpoC* R | CTTTCACGACGTACTTTAGA |
| qPCR for *sdrM* F | GCAATGATCGCAATCGGTAT |
| qPCR for *sdrM* R | GGCATAGTTGGCAGTGTTTG |
| qPCR for *lmrS* F | TGCGATGGCGATGTAGATAAA |
| qPCR for *lmrS* R | CTCACATGGCACGGCTATTA |
| qPCR for *sepA* F | CCATGATGACCCAAAAATCG |
| qPCR for *sepA* R | TTAGAGGCGCGACTTTTCAT |
| Check Amplification 1 F | TATGCCTCCTGCTGAGTTTG |
| Check Amplification 1 R | CCTTGGTCAAGCGGTTAAGA |
| Check Amplification 2,9 F | TGTATAAGACACACCACCTAAGAAA |
| Check Amplification 2 R | GCTATGTGTGGACGGGATAAG |
| Check Amplification 3 R | CTTGACTGCGAGACCTACAA |
| Check Amplification 4 F | GAAGTGCGTATTCAATGGAGAGT |
| Check Amplification 4 R | CTAGCTGTGTTGGCTTTCT |
| Check Amplification 5,6 F | ATCGCATGAAATACCTCCTTCT |
| Check Amplification 5 R | TGGGATACTACCCTAGCTGTG |
| Check Amplification 7 F | CTCGAAAGTAATTCGCCCACTA |
| Check Amplification 6,7 R | AGAAGAGCCGCAGTGAATAG |
| Check Amplification 8 F | CGTAAAGGTTCGATGTCCAAAG |
| Check Amplification 8 R | GGAGTCAGAACATGGGTGATAAG |
| Check Amplification 10 F | GCACTTTCAGGTACCTCCTTAG |
| Check Amplification 11 F | TCCATAATACGACCCTGTTGATTTA |
| Check Amplification 9,10,11,12 R | GCGACTTTCTGGTCTGTAACT |
| Check Amplification 12 F | ACCTTAAACCTTCTGCCAATCT |
| Check Amplification 3,13,14,15 F | ACGTAAATCTTCTGTTGCAGTTG |
| Check Amplification 13 R | GCGGCGTGCCTAATACAT |
| Check Amplification 14, 15 R | GCTGGATCACCTCCTTTCTAAG |
| Check Amplification 16 F | GTGTTGGGTCCCTTCGTATAAT |
| Check Amplification 16 R | TGCAATAGCCTCCGGTAAAG |

***** Underlined sequences represent homology to the respective plasmid for Gibson assembly.
